# Co-Citation Percentile Rank and JYUcite: a new network-standardized output-level citation influence metric and its implementation using Dimensions API

**DOI:** 10.1101/2020.09.23.310052

**Authors:** Janne-Tuomas Seppänen, Hanna Värri, Irene Ylönen

**Affiliations:** Open Science Centre, University of Jyväskylä, Finland

## Abstract

Judging value of scholarly outputs quantitatively remains a difficult but unavoidable challenge. Most of the proposed solutions suffer from three fundamental shortcomings: they involve i) the concept of journal, in one way or another, ii) calculating arithmetic averages from extremely skewed distributions, and iii) binning data by calendar year. Here, we introduce a new metric Co-citation Percentile Rank (CPR), that relates the current citation rate of the target output taken at resolution of days since first citable, to the distribution of current citation rates of outputs in its co-citation set, as its percentile rank in that set. We explore some of its properties with an example dataset of all scholarly outputs from University of Jyväskylä spanning multiple years and disciplines. We also demonstrate how CPR can be efficiently implemented with Dimensions database API, and provide a publicly available web resource JYUcite, allowing anyone to retrieve CPR value for any output that has a DOI and is indexed in the Dimensions database. Finally, we discuss how CPR remedies failures of the Relative Citation Ratio (RCR), and remaining issues in situations where CPR too could potentially lead to biased judgement of value.

## Introduction

Judging value of a scholarly output^1^ remains a difficult but unavoidable challenge. It is difficult because the ideals of judging scholarly value – thorough careful reading, long deliberation, debating in person or via published exchanges, peer review, replication – defeat their purpose in situations where the need for judgement in the first place arises from available time, attention and expertise resources being far exceed by the material. These situations, and hence judgement by less than ideal methods, are unavoidable: they arise in grant review panels and academic hiring committees, for science journalists, for evidence-based governance, for teachers, not to mention in daily attention allocation decisions of individual researchers.

Judging is also difficult because different aspects of value, such as *importance* and *influence*, are multifaceted concepts weighed by subjective considerations. Furthermore, influence and importance, however one measures them, are entangled but not interchangeable measures, further confounding the task. Appearing on a more prestigious platform obviously does make an output more influential, by virtue of being noticed more. Being cited more is both evidence of and path towards more influence (Wang et al 2013).

But appearance on a prestigious platform and raw citation count are not meaningful proxies for importance.

Platform prestige is a largely intangible and informally propagated social construct of academic communities. In the case of journals, it is nonetheless influenced by and correlated with the Journal Impact Factor, JIF (Garfield 1955, Garfield 2006), the annually published quantitative metric of citation performance of journals. But regardless of the degree or pretension of formality, platform prestige is always an inherently flawed proxy to judge value of individual outputs (Seglen 1997, Stringer et al 2008, Pulverer 2013, Johnston 2013). The prestige of a platform arises from high submission volumes coupled with extreme selection exclusivity, and importance of some past exceptional outputs – in other words, platform prestige mostly reflects lack of importance of a mass of material and extremity of importance of a very small number of outputs **other** than the individual output in question. It is therefore not surprising that platform prestige does not correlate positively with quality, reliability or importance of outputs published on it (Brembs 2018)

Raw citation count of an output – basis of the widely used h-index (Hirsch 2005, Norris and Oppenheim 2010) – is first and foremost predicted by the size and citation traditions of the field and obviously the age of the output (Radicchi & Castellano 2012), not the content of the output. Variation due to field of research is not limited to differences between entirely different disciplines such as Mathematics versus Medicine, but also includes significant variation between narrow sub-fields (Radicchi & Castellano 2011) and even between study questions falling within the scope of a single topical journal (Radicchi et al 2008, Opthof & Leydesdorff 2010). Annual global publication volumes and citation patterns such as typical length of reference lists and typical age of cited publications can be very different between, say, Theoretical Ecology versus Environmental Ecology, or between studies investigating genetic basis of behaviour in *Orcinus orca* versus in *Caenorhabditis elegans*, both published on some same Behavioural Ecology platform.

These issues have been recognised and debated for a long time. Proposed solutions include radical reassessment of the value of “value” in scholarly work, ranging from fundamental sociological and linguistic changes to abandon “fetishisation of excellence” (Moore et al 2017), to replacing research grant review panels with random lotteries (Fang 2016) as was recently done by The Health Research Council of New Zealand (Liu et al. 2020). These approaches merit serious consideration and real-world testing and adoption, but are outside the scope of this article.

Others recognise the usefulness of bibliometrics, or at least their unavoidability, as long as they are better than the obviously maligned things of yore. These solutions tend to be new metrics, and there are hundreds of them. Wildgaard (2014) and Waltmann (2016) provide recent reviews. A non-exhaustive list includes: normalizing citation counts by averages of articles appearing in a single journal or in journal categories assigned by publisher or database vendor (Opthof & Leydesdorff 2010, Waltman et al 2011a, 2011b, Bornmann & Leydesdorff 2013); normalizing citation counts by average number of cited references in citing articles (Zitt & Small 2008, Moed 2010); counting number of articles in the “top 10%” of most cited articles but normalizing by researcher’s (academic) age (Bornmann & Marx 2013); citation network centrality measures such as the journal Eigenfactor and the derivative Article Influence which is journal Eigenfactor divided by the number of articles on that platform (as Bergström 2007 admits, this is essentially judging “importance of an article by the company it keeps” as thus no better than the misuse of JIF to judge individual articles); dealing with collaborative authorship by fractioning the credit among individual researchers (e.g. Schreiber 2009), their institutions (Perianes-Rodríguez & Ruiz-Castillo 2015) or countries (Aksnes et al 2012).

Most of the proposed solutions suffer from three fundamental shortcomings:

1. They involve the concept of journal, in one way or another.
2. They involve calculating arithmetic averages from extremely skewed distributions.
3. They involve binning data by calendar year.

All of these probably first emerged historically from only aggregate and annually compiled data being easily available in the pre-internet era, and later became staple traditions seen to hold some value in themselves. But today they just bring unnecessary inaccuracy, bias, restricted samples, delays and lack of generality to measures seeking to judge value of individual scholarly outputs and researchers.

## Methods

### Co-citation Percentile Rank (CPR)

Here, we describe Co-citation Percentile Rank (CPR), a metric that relates the current citation rate of the target output taken at resolution of days since first citable, to the distribution of current citation rates of outputs in its co-citation set (Fig 1.), as its percentile rank in that set. The co-citation set is defined identically to “co-citation network” in Hutchins et al (2016), see Fig 1. and Discussion.

**Fig 1.**
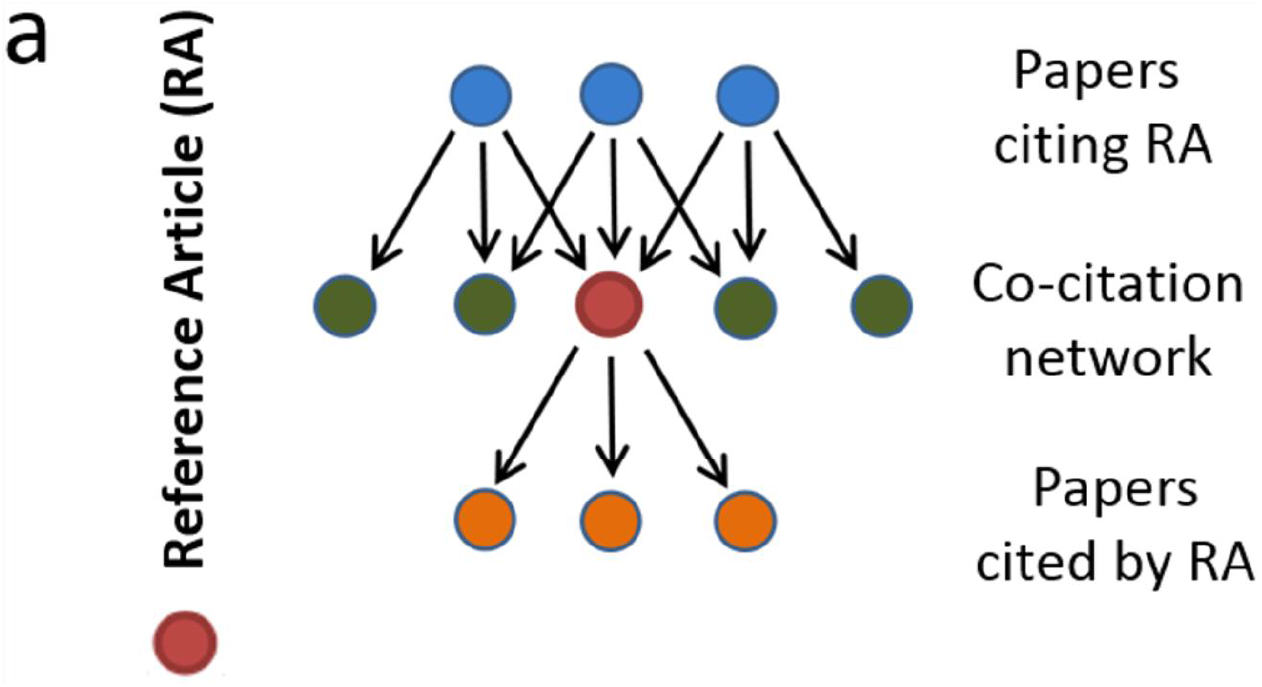
Adopted from Hutchins et al (2016) “Fig 1a. Schematic of a co-citation network. The reference article (RA) (red, middle row) cites previous papers from the literature (orange, bottom row); subsequent papers cite the RA (blue, top row). The co-citation network is the set of papers that appear alongside the article in the subsequent citing papers (green, middle row). The field citation rate is calculated as the mean of the latter articles’ journal citation rates.”

To retrieve the co-citation set and its bibliometric metadata, we use Dimensions Search Language (DSL) on their API:

1. get Dimensions **ID** and other data about the target, by **DOI** (or multiple targets: up to 400 can be queried in a single call, then looped through the queries below): search publications where doi in [**DOI**] return publications [basics + extras + date_inserted]
2. get data about outputs where the target appears in their list of references, i.e. the citing outputs, and in particular the reference_ids of outputs they cite, i.e. the co-citation set of the target: search publications where reference_ids=“**ID**” return publications [title+id+reference_ids]
3. Collect the reference_ids from the result array, remove duplicates, concatenate the unique ids into comma-separated strings chunked to max 400 ids per string.
4. get metadata about the co-citation set, by **reference_ids** string (looping through if multiple chunks): search publications where id in [“**reference_ids** “] return publications [title + doi + times_cited + date + date_inserted + id]

Once the citation counts and days since the output became citable have been assembled, these are converted to 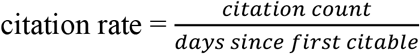. When used in graphs and tables, that citation rate is multiplied by 365 to get *“citations /365 days since becoming citable”*. We do this to aid interpretation and comparisons, but we emphasize that 365 days is not a calendar year, and that the citation rate is defined also for outputs that are less than one year old.

With data as extremely skewed as citation counts, it should be obvious that arithmetic averages are not appropriate denominators. Leydesdorff et al (2011) make this case explicitly for bibliometrics, and argue that newly proposed metrics should use percentile ranks instead of ratios to arithmetic averages of reference sets.

The co-citation set is then ordered by citation rate, and then

### CPR = percentage of citation rates in the co-citation set that are less than or equal to the citation rate of the target output

CPR thus ranges from 0 (= all outputs in the co-citation set are cited more frequently than the target) to 100 (= none of the outputs in the co-citation set are cited more frequently than the target). CPR is undefined for targets that have not been cited, or for which the algorithm is unable to find co-citation set metadata. The quartiles and average of the citation rates, and the size of the co-citation set are also calculated and saved for illustrative purpose.

Note on the day count used in the algorithm: when an output first becomes citable, typically as an “early online” article, Dimensions lists that date in the metadata. However, for platforms that later assign outputs to issues (some even distribute those arbitrary collections of outputs as anachronistic physical representations created on cellulose-based perishable material), that date eventually gets replaced in Dimensions metadata by a later so-called “publication date”. The original, true date when the work first became citable, is not currently retained in Dimensions in these cases. For some platforms the resulting error may be just few tens of days, but in others the error may be many hundreds of days. In most cases the true date would be recoverable from metadata in CrossRef or the landing page of the output DOI, but this would impose prohibitive time performance cost on the algorithm. We hope this unfortunate and unnecessary source of error gets remedied in the future. For now, we resort to retrieving also the **date_inserted** value alongside the **date** value from Dimensions API, and choose the earlier of these two dates, acknowledging some outputs may still enter the algorithm with unknown error in their citation rate, and that this error is more common and likely larger for some disciplines, such as Economics or Humanities.

The source code of the algorithm is publicly available available at https://gitlab.jyu.fi/jyucite/published_cpr and published in Seppänen (2020)

### Example dataset

Metadata for total of 41 713 outputs from JYU current research system published between 2007-2019 (all kinds, including non-peer-reviewed outputs) were assembled and evaluated, out of which 13 337 i) were discoverable in Dimensions by either DOI or title and ii) had at least one citer.

That dataset of metadata for 13 337 outputs is published alongside this article. It gives DOI, Dimensions id, title, type of output (as defined by Ministry of Education and Culture in Finland (2019), and assigned by information specialists at research organizations), university home department(s) at JYU, publication date, number of days available and number of citations accrued up to date of analysis, calculated citation rate per 365 days, co-citation set size, number of co-citations for which times_cited metadata was present, co-citation median citation rate, quartiles and average, the CPR metric, and the solve time (seconds it took to retrieve the co-citation set and calculate the metrics) as well as UNIX timestamp of the time when the calculation was done.

Out of those, 13 170 had at least 10 co-citations for which metadata could be found, and these were retained for beta regression analysis.

The example dataset is is publicly available at https://gitlab.jyu.fi/jyucite/published_cpr and published in Seppänen (2020)

### Statistical analysis

Descriptive analyses on the effect of number of citers on CPR solving time and size of the co-citation set were done with quantile regression (Koenker 2020). Quantile regression model fits are estimated using the R1 statistic (Koenker & Machado 1999, Long 2020).

Because the distribution of CPR data is by definition bound to unit interval (0,100), and, as is typical for any citation metric, also heteroscedastic and asymmetric, we model its response to predictors using beta regression (Cribari-Neto & Zeileis 2010). Analysis incorporating precision parameter to account for heteroscedasticity failed to converge with raw data for any link function, so a preliminary analysis was run without precision parameter to identify outliers. Model with log-log link function converged so that is used in all subsequent analyses. Twenty extremely highly cited outputs showed outlier impact (Cook’s distance > 1.0) in preliminary regression analysis and were removed from subsequent analysis. Beta regression including a precision parameter was then run. Cases where CPR was zero (405 cases) continued to have disproportionate generalized leverage on the fit, and were also removed. The final sample where both extreme highs and lows had thus been removed contained 12740 observations.

After fitting the overall beta regression of CPR to citation rate, we next explore the differences in that relationship between academic disciplines. Each output in the database is affiliated with one or more academic departments at University of Jyväskylä (JYU). For purposes of this analysis, we simplify the data by merging data from units falling under same discipline (e.g. Institute for Education Research is merged with Department of Education). For outputs still having multiple affiliations after the mergers, the record is replicated so that an output occurs once for each affiliated department. Departments having fewer than 100 outputs were excluded from the final analysis. The derived expanded dataset contains 13871 observations from 16 different academic departments (Table 1.)

**Table 1.** Departments and output counts included in the analysis.

| *Department* | *No. outputs* |
| --- | --- |
| Physics | 2469 |
| Sport & Health | 2117 |
| Bio-environmental | 1912 |
| Information Technology | 1475 |
| Chemistry | 1415 |
| Psychology | 1047 |
| Education | 586 |
| Mathematics & Statistics | 585 |
| Business & Economics | 499 |
| Social Sciences & Philosophy | 442 |
| Language & Communication | 349 |
| Teacher Education | 312 |
| Music, Arts & Culture | 273 |
| Agora Center* | 185 |
| History & Ethnology | 104 |
| Applied Linguistics | 101 |

\*multidisciplinary institute on humanities, social sciences and technology, active 2012-2017

We then model CPR as a function of citation rate and academic department, including interaction terms and a precision parameter accounting for heteroscedasticity along the citation rate range.

The R code to replicate the analyses and figures presented here is publicly available at https://gitlab.jyu.fi/jyucite/published_cpr and published in Seppänen (2020)

## Results

With our computing setup (a single RHEL7.7 Server, 1 x 2.60GHz CPU, 2GB RAM, PHP 5.4.16, in university’s fiber-optic network), solving CPR for an output was within reasonable time performance and solve time was linearly scaled with number of citers, though variance was considerable (Fig. 2).

**Fig. 2.**
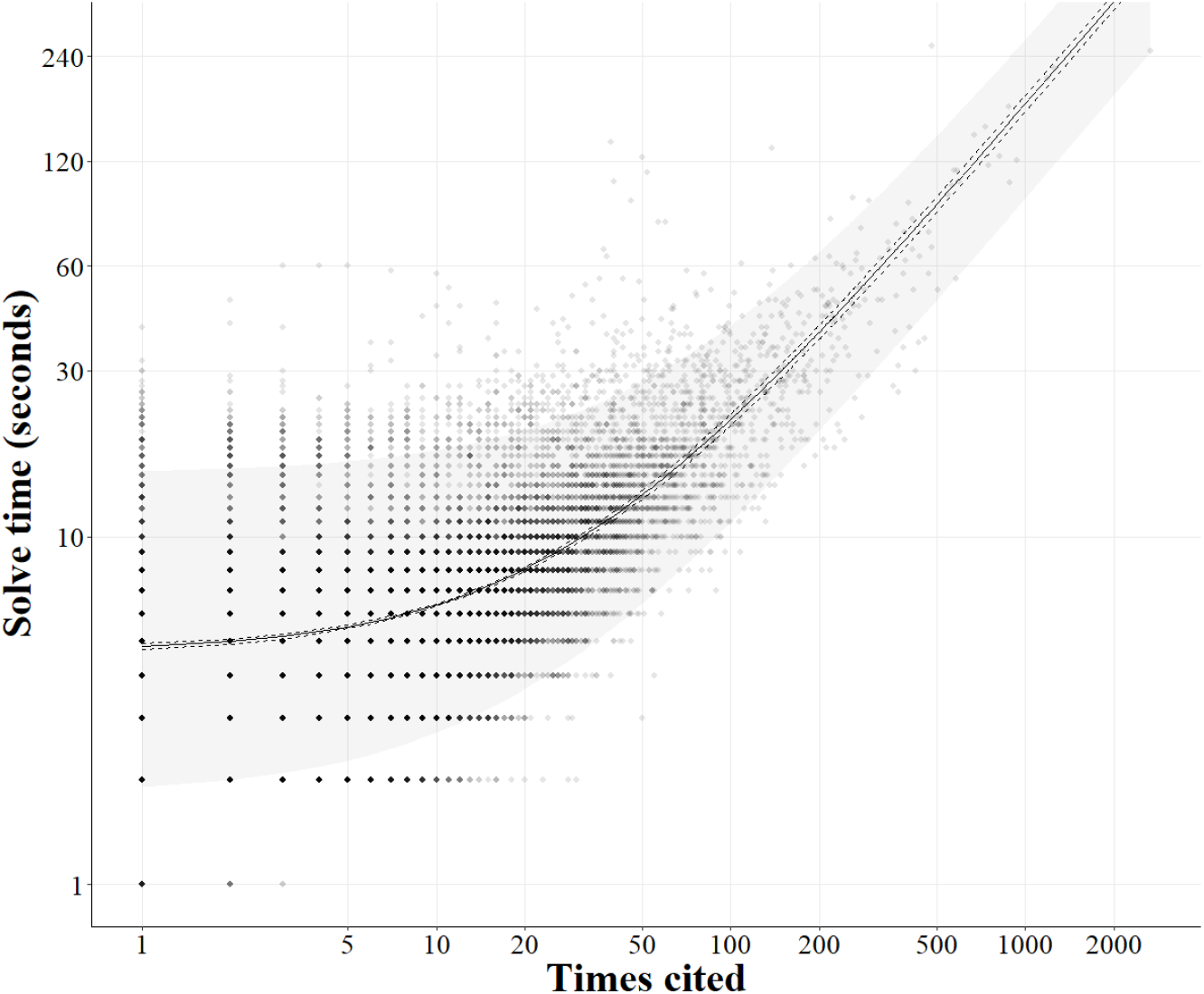
Log-log plot of the CPR solve time as a function of times the target output is cited. Dot opacity is set at 0.1 in R plotting function so that overlap darkness illustrates density of observations. Quantile regression for median (line) and 90% prediction interval (shaded area) and 99% confidence interval (dashed lines). Solve time ~ 0.17 * times cited + 4.66 (N = 13 337, model fit: R^1^(0.5) = 0.29). The variation in solve time results from variable co-citation set size given times cited (see Fig 3.), and occasional automatic delays in the algorithm as it adheres to rate limit of max 30 requests per minute as mandated in Dimensions’s terms of use.

Co-citation set size expanded rapidly as number of citers grew: each new citer typically brought ca. 40 new entries to output’s co-citation set, and occasional review, book or reference work citer could bring hundreds or even thousands of entries (Fig. 3) Overall beta regression of CPR as a function of citation rate illustrates that though CPR increases with increasing citation rate, the relationship has considerable variation, as should be expected if CPR captures sub-field specific citation influence. Some outputs achieve CPR around 50 (i.e. are cited at least as frequently as half of their co-citations) when they get cited 1-2 times per 365 days, while other outputs need 9-10 citations per 365 days to achieve similar CPR (Fig 4.).

**Fig. 3.**
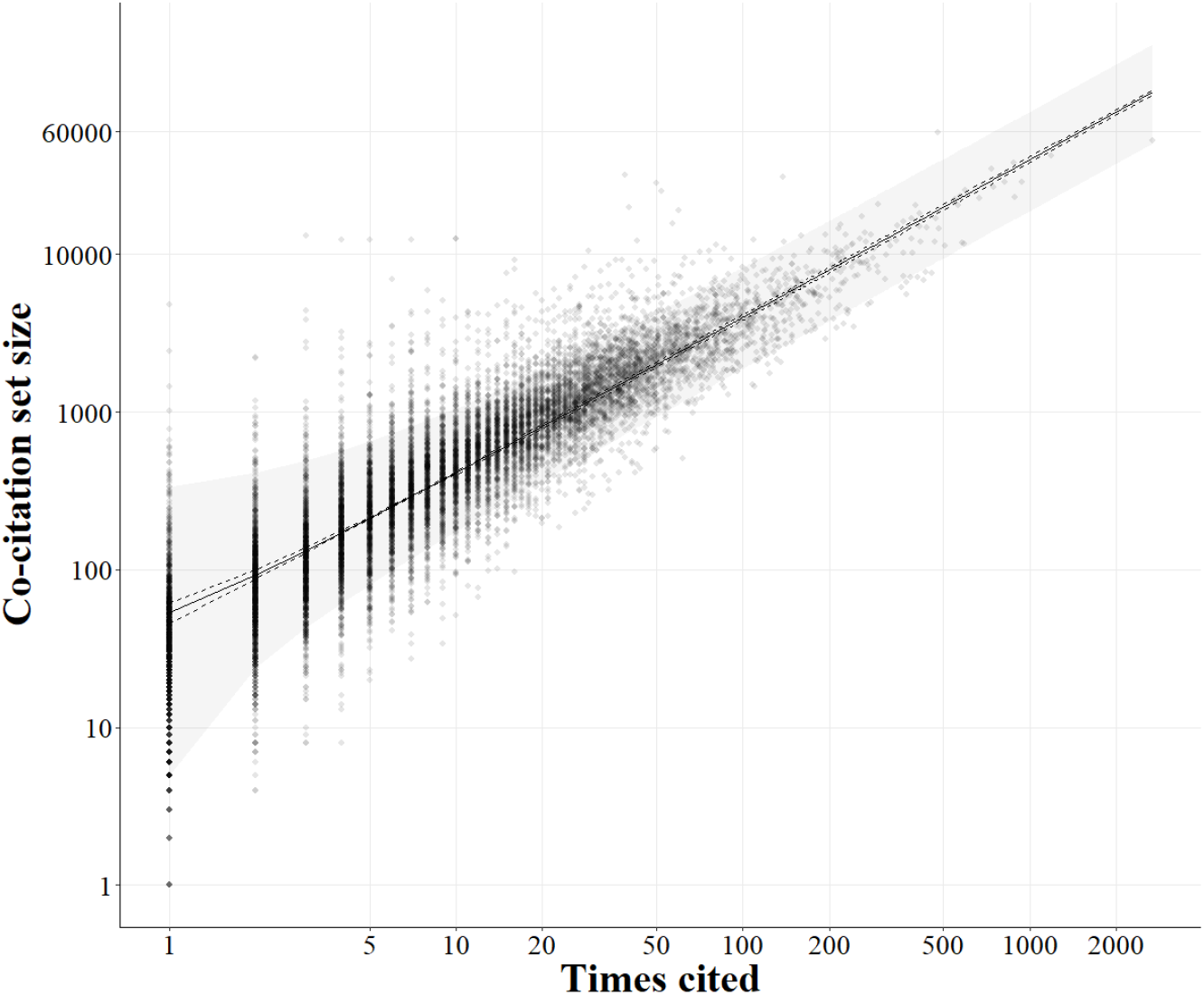
Log-log plot of the co-citation set size as function of times a peer-reviewed output has been cited. Dot opacity is set at 0.1 in R plotting function so that overlap darkness illustrates density of observations. Quantile regression for median (line) and 90% prediction interval (shaded area) and 99% confidence interval (dashed lines). Co-citation set size ~ 39.73 * times cited + 13.82 (N = 13337, model fit: R^1^(0.5) = 0.54). The extreme high outliers result from an output being cited by reviews and books, or large reference works, which alone can bring hundreds or thousands of entries, respectively, to output’s co-citation set.

**Fig 4.**
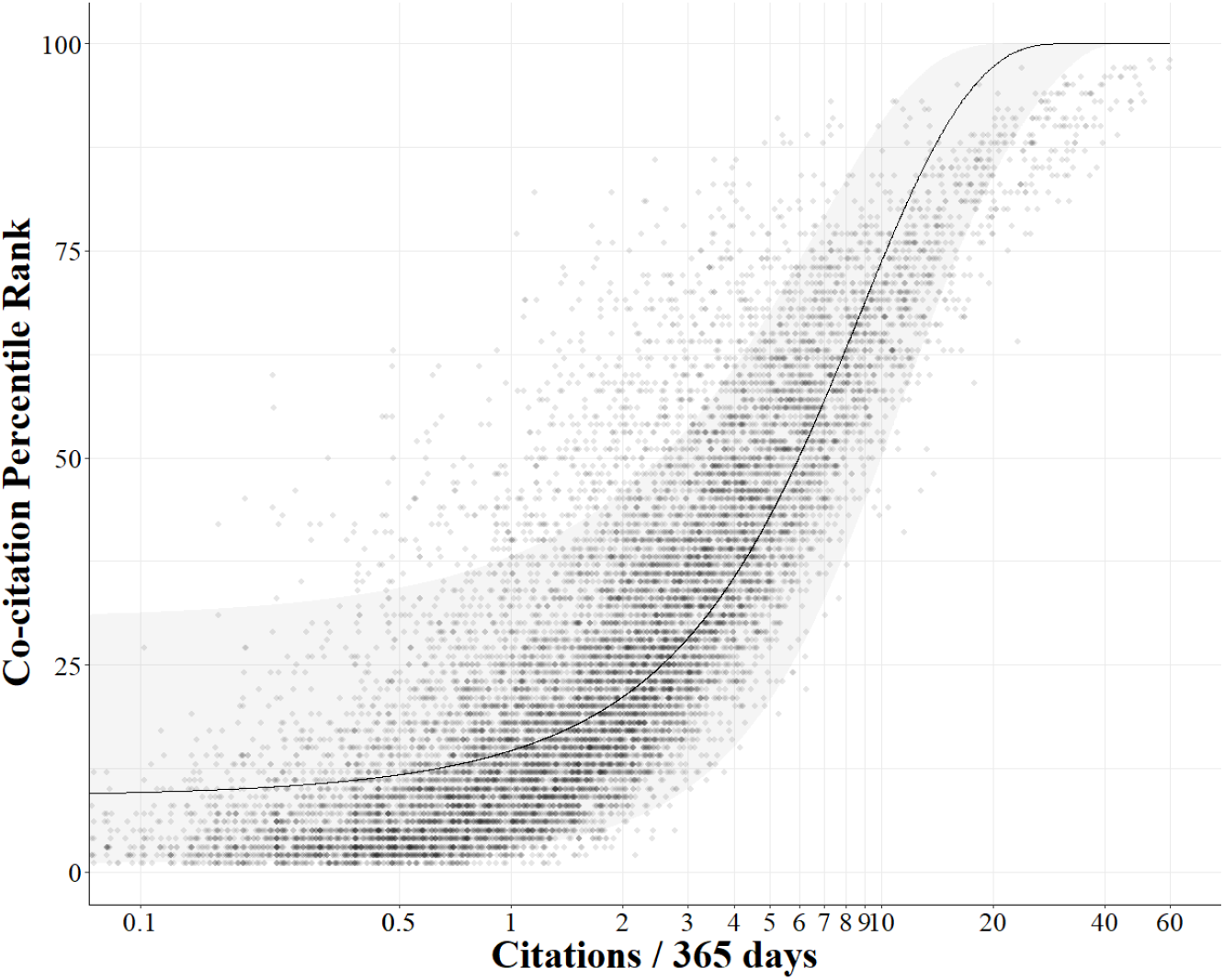
Co-citation Percentile Rank of outputs as a function of citation rate (citations / 365 days since becoming citable). Dot opacity is set at 0.1 in R plotting function so that overlap darkness illustrates density of observations. Beta regression predicted median (line) and 90% prediction interval (shaded area).

With the expanded derived data, likelihood ratio test shows that including departments in the regression model gives significantly better fit (*χ*^2^ = 1331.3, df = 30, p < 0.001), despite being considerably more complex model (34 degrees of freedom vs 4 when modelling as function of citation rate only: ΔAIC = 1271.3). In other words, some difference between academic departments explains significant part of the variability in response of CPR to citation rate.

Different global publication volumes and citation behaviours in different disciplines are the obvious likely explanation.

Pairwise post-hoc Tukey tests (Supplementary Table S1) show that majority of the department contrasts in coefficient for citation rate on CPR (69 out of 120) are statistically significant (Bonferroni-adjusted p-value < 0.05). Differences are most pronounced between Mathematics & Statistics vs other disciplines: the slope is consistently steeper in Mathematics & Statistics, i.e. CPR responds more rapidly to increase in citation rate. Going from rate of two citations to five citations per 365 days is predicted to move an output in Mathematics & Statistics from bottom third to top third in citation rate in it’s co-citation set. In contrast at the other extreme, in Physics same change in citation rate means an output moves from bottom 25% only to a rank still below median in its co-citation set.

Similarly, post-hoc tests for contrasts in intercept term (Supplementary Table S2) show that majority of the department contrasts (65 out of 120) are statistically significant. The Agora Center (a former multidisciplinary research unit bringing together humanities, social sciences and information technology) differs from all other department by having significantly lower intercept, i.e. CPR of Agora Center outputs was generally lower, given a citation rate. This could partially be a result of the multidisciplinary nature the unit had (see section 4.1. below). History & Ethnology on the other hand consistently has significantly higher intercept term then other departments, i.e. CPR for a History & Ethnology output tends to be higher, given a certain citation rate.

**Fig 5.**
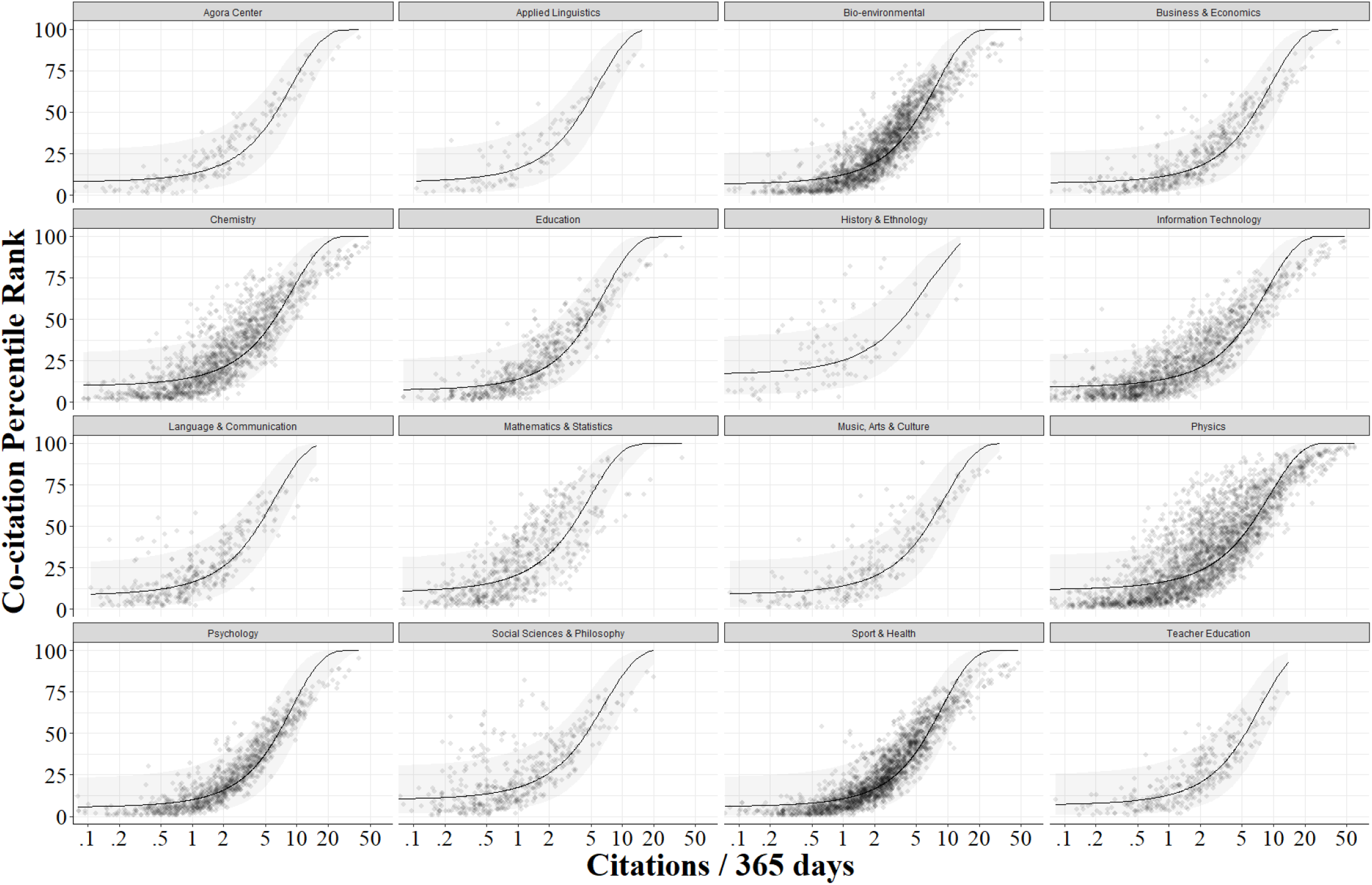
Co-citation Percentile Rank of outputs as a function of citation rate (citations/ 365 days since becoming citable) for 16 different academic departments at JYU. Dot opacity is set at 0.1 in R plotting function so that overlap darkness illustrates density of observations. Beta regression predicted mean (line) and 90% prediction interval (shaded area). Note that x-axis is logarithmic.

## Discussion

### CPR reflects expected differences between disciplines

The contrast between academic disciplines found are consistent with conventional wisdom and highlight the utility of CPR as a field-normalized citation rate metric. In Mathematics & Statistics, publication volumes are relatively low both individually and collectively, and works often cite just a few foundational references. In Physics, the three particle accelerators and nanoscale materials research at JYU result in outputs that are in very large, fast-moving fields where massively collaborative works get cited quickly. In History & Ethnology, the discipline has smaller and slower publication volumes as many outputs are monographs, and perhaps citations behaviours are also more siloed to fine-grained sub-fields and by language. It should be noted though, that the sample size for History and Ethnology is small here, and furthermore that sparse sample shows more variation than other disciplines (Fig 4.), so inference must be cautious

### Implemetation using Dimensions API

The efficiency of the Dimensions API is remarkable for this task: for an ouvre of M target outputs (say, the publication list of a person), only 1+M queries are needed to obtain the IDs of entire co-citation sets of all target outputs. The N IDs in a single co-citation set can often number thousands. But metadata for these can be retrieved in N/400 batch queries. Because the algorithm does not have to make separate Dimensions API queries to retrieve either the co-citation set IDs or the metadata for all of the entries, the CPR solve time is reasonably linearly scaled to the number of citers citing the target output (using our computer and network resources, ca. 0.17 seconds / citer, Fig 2.). Performance could be in principle multiplied by parallelization, but only with consent of the database provider. Here, we operated under the standard consumer terms of service by Dimensions, which set a maximum request rate of 30 / minute.

Interestingly, whereas Hutchins et al (2016) found that each citer added on average ca. 18 new outputs to the co-citation set, we find that each new citer adds ca. 40 (Fig 3.). This may result from the narrower focus as Hutchins et al limited their analysis to articles in those journals in which NIH R01-funded researchers published between 2002–2012, i.e. largely biomedical research. In contrast, the dataset we use here to illustrate CPR covers all research outputs 2007-2019 from one university, spanning 18 different departments. Second, and likely a more consequential difference is that Hutchins et al used Web of Science, whereas we use Dimensions citation database. Dimensions indexes citations much more inclusively, particularly in dissertations, textbooks, monographs and large reference works such as encyclopaedias. Such book-length sources often include much longer lists of references – and hence large contributions to co-citation sets – than typical articles.

Faster accrual of the co-citation set using Dimensions also means our implementation of CPR is not very vulnerable to finite number effects. For median, typical target output, mere three citers bring its co-citation set size above 100, where percentile can be confidently stated as integer value without having to interpolate between observed values.

### CPR and RCR

Definition of the reference set, and inspiration for developing CPR, is the Relative Citation Ratio (RCR, Hutchins et al 2016). The foundational idea of RCR – comparing the citation rate of target output to those of ‘peer’ outputs which are cited alongside the target, the co-citation set – makes intuitive sense and is a truly journal-independent research field – normalization for quantitative comparison of academic impact.

All three sets of outputs in an output’s citation network (cited, citing and co-citations, see Fig 1.) can be intuitively seen to “define empirical field boundaries that are simultaneously more flexible and more precise” (Hutchins et al 2016). But the co-citation set is clearly the superior choice. While the cited set is defined just by the authors, the other two sets grow dynamically over time and can be seen to reflect current aggregate expert opinion of what constitutes the field of the target output. The citing set might be considered to be the most direct expression of that aggregate expert opinion, but as a reference set for comparing citation performance, the set suffers from two significant problems: by definition it only includes newer outputs than the target, and it usually grows slowly. Hence, typically for years there would not be a meaningful comparison set. Then a set would begin to exist but comparison would be meaningless as the citing outputs would not yet have had time to get cited themselves. And if and once, eventually, those issues begin to fade then the target output gets compared to a set that contains more currently relevant and derived research, only, several years after the target itself was at its most influential. In slow-moving fields, this could easily take a decade or longer.

In contrast, the co-citation set grows by multiple outputs every time the target is cited. Even with partial overlap, Hutchins et al (2016) found that each citer added on average ca. 18 new outputs to the co-citation set in their example in biomedical sciences. Just as importantly, the temporal coverage of the co-citation set encompasses all research from oldest seminal outputs to the newest research that can reasonably appear in reference lists today.

In addition, the text similarity (Lewis et al 2006) was greater among co-cited outputs than among articles appearing on same platform.

However, RCR is problematic in several ways, and CPR has been developed to remedy those failures.

Most glaring discrepancy to the promise of the idea of truly output-level metric, was that Hutchins et al resorted to using Journal Impact Factors (JIF) in the normalization. They define Field Citation Rate (FCR) as the average JIF of journals where the co-cited articles appeared. The only justification for this decision was a vague claim that average of citation rates of articles would be “vulnerable to finite number effects”. An unmentioned but possible explanation was to make the introduction presented in their scholarly article (based on Web of Science citation data) comparable with the online implementation (based on PubMed data). PubMed indexes an extremely narrow scope of the published literature and even narrower slice of the citation network, so most articles in the co-citation set would show zero or very few citations. Such zero-inflated and relatively small data indeed would be susceptible to instability of average, but that is more a property of the data source, rather than the co-citation set itself.

Regardless of justification, JIF itself is (sort of) an average, from an extremely skewed distribution. Furthermore, calculation of JIF is a negotiated and extensively gamed process in the publishing industry, and most importantly, it is trivially obvious that neither the true importance nor the citation rates of individual outputs are meaningfully correlated with the JIF of the journal where the output appears.

The target Article Citation Rate (ACR) was defined as the accrued citations divided by the number of calendar years since publishing, excluding the publication year. This introduces another serious yet unnecessary problem: some articles had had 364 days more to accrue citations than others divided by the same number of calendar years. Especially for recent articles this is a significant source of error. Modern citation databases are updated continuously and outputs and their citations appear within days, sometimes before, of the date of publication, and the date is listed in the metadata. There is no reason to obscure that temporal resolution of citation data by aggregating counts by calendar years, when citation rates can be easily expressed as citations / days since citable.

Then, instead of straightforward definition of RCR = ACR/FCR, Hutchings et al (2016) present a convoluted regression normalization procedure requiring very large datasets of other “benchmarking” articles. This decision may reflect an unmentioned recognition that their FCR, as it is an average from an extremely skewed distribution of JIF values which themselves are (sort of) averages from extremely skewed distribution of two-year article citation counts, inevitably is a much larger number than the expected citation rate of a typical article. As Janssens et al (2017) notes, the FCR-to-RCR regressions had intercepts ranging from 0.12 to 1.95 and slopes from 0.50 to 0.78, which is entirely expected given the way FCR is calculated. The regression normalization procedure yielded Expected Citation Rate (ECR, detailed in Hutchins et al 2016b) for the target article, and then RCR = ACR/ECR so that for articles in the dataset with the same FCR, the article with average citation rate had RCR = 1.0. Again, given the skewed nature of citation accrual, this inevitably means that most articles will have an RCR (much) below 1.0 despite many of them being cited more frequently than majority of articles in their co-citation sets.

Final shortcoming was the subsequent implementation of RCR in iCite, building the algorithm on top of PubMed citation data. That data is woefully sparse for most fields of science. Most outputs simply are not indexed by PubMed at all. And even when they are indexed, PubMed fails to capture vast majority of citations to them.

### JYUcite

We have built a publicly available web resource allowing anyone to retrieve CPR value for any output that has a DOI and is indexed in the Dimensions database. The resource is available at https://oscsolutions.cc.jyu.fi/jyucite

Additionally, JYUcite graphically and numerically displays the observed citation rate, and size N and quartiles of citation rates in the co-citation set (Fig 5.). The citation rates are expressed rescaled as citations / 365 days (note: not a calendar year, but daily rate scaled to one year for convenience – see Methods) At the time of publication of this article, JYUcite limits number of DOIs per request to 10 to ensure performance as we develop the resource further. JYUcite also limits the number of newly calculated CPR values returned (i.e. those requiring JYUcite server to make calls to Dimensions API) to 100 per IP-address per day, as contractually agreed with Dimensions database. If a CPR value for a DOI has already been calculated previously and is saved in JYUcite’s own database and is younger than 100 days, it is returned immediately without new calculation and does not count in the daily rate limit.

**Fig 5.**
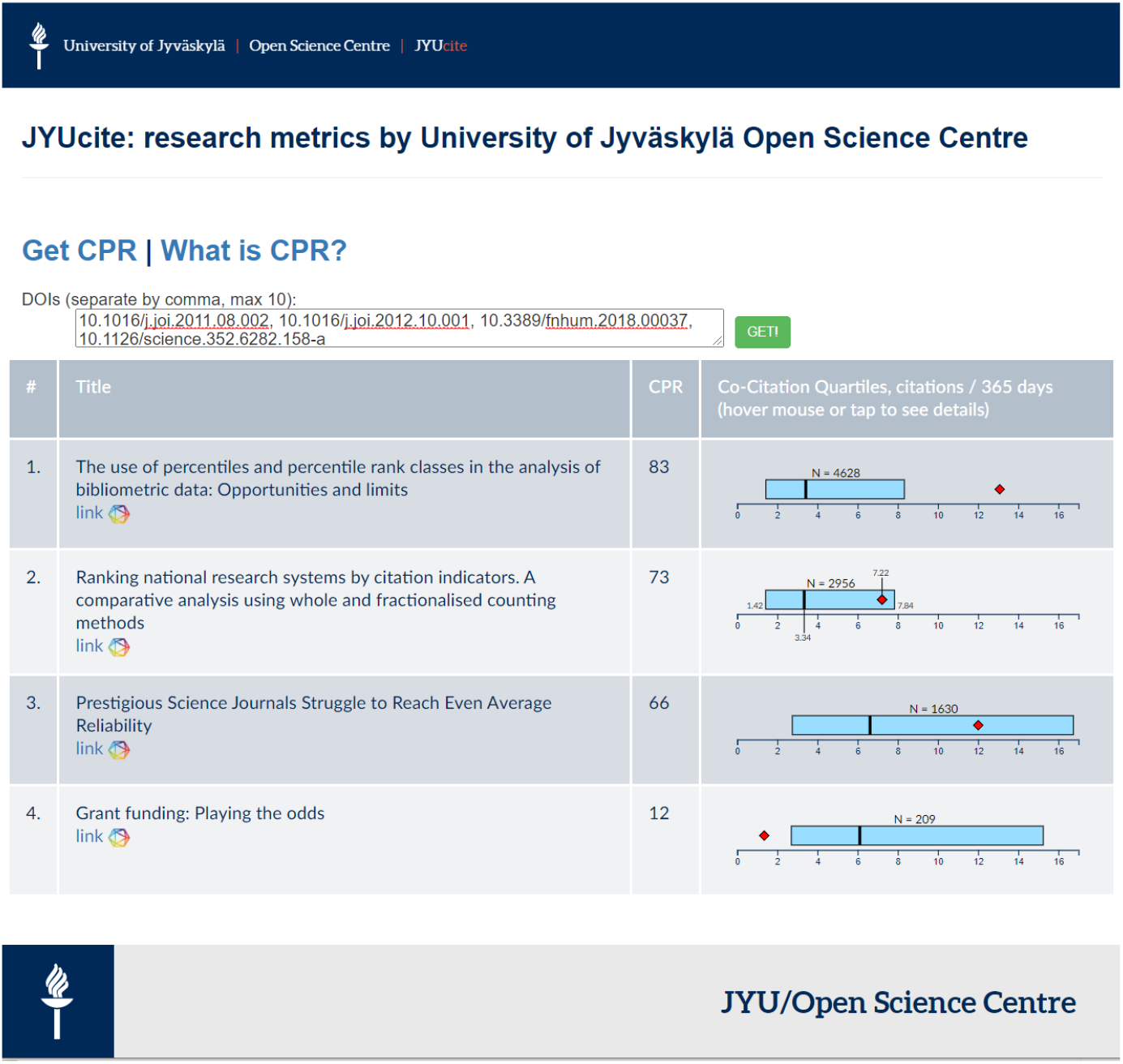
Screenshot from JYUcite web service, showing results view when requesting CPR for some articles listed in References in this article.

### Remaining issues

#### Interdisciplinary influence

An output may suffer lowering CPR if it begins to get cited in another field where outputs typically get much more citations (e.g. due to field size and typical lengths of reference lists). It would begin to gain (proportionally large numbers if the other field typically has longer length of reference lists) relatively frequently cited co-citations into its co-citation set, repressing its CPR. Thus interdisciplinary influence, which typically is seen as a merit, may actually erode the CPR value of an output (see also Waltman 2015). On the other hand, it could also be argued that once an output begins to become relevant in another discipline, it **should** be start to get compared to outputs in that field, otherwise it would appear unduly influential compared to those.

A possible remedy could be to pay attention to relative 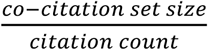 ratio: if an output in e.g. individual researcher’s publication list stands out from others with a noticeably larger ratio, it might indicate interdisciplinary influence, and that output with its citing set should then be flagged for individual judgement by a human, for potential extra merit.

The reverse (an output from a field with high typical citation rates begins to get cited in a field with typically much lower citation rates, thereby gaining low-cited co-citations, inflating its CPR) is likely to have less significant impact as the gain of low-cited co-citations is slow compared to gain of co-citations in its original field, and the proportion of those co-citations remains marginal in the set.

#### Co-citations in reference works

Being cited in a large reference work, such as an encyclopaedia, can bring thousands of entries into the co-citation set. On one hand this is an advantage, e.g. when “Encyclopedia of Library and Information Sciences” creates relevant co-citation set of 4608 entries for large number of outputs, which otherwise would wait many years to accrue similar sized comparison set.

However, “Encyclopedia of Library and Information Sciences” also cites articles from completely unrelated fields, e.g. a conservation ecology article from 1968 titled “Chlorinated Hydrocarbons and Eggshell Changes in Raptorial and Fish-Eating Birds”. While this is likely to be an insignificant source of bias for the CPR of outputs in Library and Information Sciences (relatively few of the thousands of co-citations accrued are from outside fields), it could be significant for the CPR of the out-field article cited. It seems likely though, that outputs that end up cited in reference works outside their own field are cited extensively in their old fields before that, and tend to be aged. The example article above has, at time of writing this, been cited 301 times and has a co-citation set of 21.616 outputs, so contribution of the encyclopaedia is at most 20% of the co-citation set.

A possible remedy could be to exclude co-citations coming from citers that have extreme outlier count of references. A typical scholarly article has ca 30 references, so a threshold of 300 would likely retain all original research and most review articles, while excluding large reference works. However, the trade-off of such filter would be to lose relevant co-citations for many little-cited articles, which could be an issue in disciplines and fields where co-citation sets accrue slowly.

#### Hyper-cited co-citations

Some scholarly outputs, particularly in natural sciences, medicine and engineering, may receive many thousands of citations every year, and consequently are likely to appear in the co-citation set of most outputs. These hyper-cited outputs tend to be statistical analysis textbooks (e.g. “Statistical Power Analysis for the Behavioral Sciences (2015)”, cited more than 37.000 times in total, or over 5000 times per every 365 days) or companion articles to statistical packages in the R-software (e.g. “Fitting Linear Mixed-Effects Models Using lme4 (2015)”, cited more than 21.000 times in total, or almost 4000 times per every 356 days) or descriptions of standard hardware or software infrastructure (e.g. “A short history of SHELX (2008)”, cited more than 68.000 times in total, or over 5000 times per every 365 days).

Is inclusion of such “boilerplate” general methodology outputs in the comparison set a source of bias? They are always far more frequently cited than a typical target output, and thus outrank them, lowering the target output’s CPR. On the other hand, they often objectively **do** have more significant influence on research than any field-specific research output. Also, they cannot be automatically excluded using either blacklists or citation rate thresholds, as the target output may well be directly in the same field as, or a co-citation have citation rate rivalling, e.g. a statistical methodology “boilerplate” reference.

But even where hyper-cited co-citations are seen as a source of bias for CPR, the magnitude of such bias decreases rapidly as target output gains more citers and thereby median of 40 new co-citations per new citer (Fig 3.). Single or few hyper-cited co-citations in a large set do not have a large effect on the percentile rank, while they would easily distort a metric relating the target to averages in the set.

#### Delayed recognition and instant fading

The value of a scholarly output is not necessarily present or recognized immediately upon publication. Some other, later discovery may suddenly reveal an important aspect of an earlier output, or its relevance in another field of research may be get re-discovered serendipitously, awakening the output to suddenly increased influence, many years after first becoming citable. Ke et al (2015) present analysis of temporal patterns for such “sleeping beauties”, and find that while there are cases of very pronounced delayed recognition and they are often associated with delayed accrual of interdisciplinary citations, such articles are not exceptional rare outliers in their own category, but simply extreme cases in heterogenous but continuous distribution.

Conversely, the influence of a scholarly output may be immediate and intense but then quickly fades (Ye & Bornmann 2018). This can happen in very rapidly advancing fields where an important discovery can spur quick succession of derivative further discoveries or improvements which overshadow their ancestors, or for big or controversial claims that generate large initial interest which however proves the claim false and the output is thereafter ignored and forgotten.

Consider three outputs with similar citation rates, where one is an old instant fade, another equally old but recently awakened sleeping beauty, and third a new output receiving few but immediate citations. Their CPRs would be similar. Whether this would be an unfair similarity is not necessarily clear: should an awakened sleeping beauty be considered more valuable than a new discovery, or should we acknowledge that CPR reflects whatever it was that kept the sleeping beauty from gaining influence sooner? Notably, the direction of change in CPR would be different for these three examples.

## Acknowledgements

We thank colleagues at Open Science Centre at JYU for enabling assembly of the dataset used in this article, and Jyrki Laitinen and colleagues at Digital Services at JYU for expertise and tireless help in setting up servers and keeping them running. Stacy Konkiel and other experts and customer support staff at Digital Science have provided valuable and timely guidance.

## Conflict of interest statement

Access to Dimensions API was obtained by otherwise standard paid subscription contract between its owner Digital Science Ltd and JYU, but special provisions were negotiated and included in that contract to allow public display of data and derived metrics at JYUcite, and to prominently display Dimensions logo and name in that web service. No provisions were made regarding academic outputs and none of the authors are affiliated or compensated by Digital Science Ltd; this work is independent. Experts employed by Digital Science Ltd may provide public peer review comments on this work where their name and affiliation is disclosed, just like any other experts.

## Source code and data availability

The source code of the algorithm, the example data, and the R code to reproduce the statistical analysis and figures here. is publicly available available at https://gitlab.jyu.fi/jyucite/published_cpr and published in Seppänen (2020)

---

**Supplementary Table S1.** Pairwise Tukey post-hoc test for difference between academic departments at JYU, in coefficient for citation rate on CPR, in decreasing order of absolute z-value. Bonferroni-adjusted p-values reported.

| **Linear Hypotheses: citation rate coefficient for CPR is equal between two departments** | **Estimate** | **Std. Error** | **z value** | **p** |  |
| --- | --- | --- | --- | --- | --- |
| Physics - Mathematics & Statistics = 0 | -0.15631 | 0.010982 | -14.233 | <0.01 | *** |
| Mathematics & Statistics - Chemistry = 0 | 0.149505 | 0.01128 | 13.255 | <0.01 | *** |
| Mathematics & Statistics - Information Technology = 0 | 0.137755 | 0.011369 | 12.117 | <0.01 | *** |
| Mathematics & Statistics - Business & Economics = 0 | 0.150434 | 0.012498 | 12.037 | <0.01 | *** |
| Sport & Health - Mathematics & Statistics = 0 | -0.13306 | 0.011177 | -11.904 | <0.01 | *** |
| Psychology - Mathematics & Statistics = 0 | -0.13685 | 0.011604 | -11.794 | <0.01 | *** |
| Music, Arts & Culture - Mathematics & Statistics = 0 | -0.15491 | 0.013349 | -11.605 | <0.01 | *** |
| Physics - Bio-environmental = 0 | -0.04895 | 0.00443 | -11.049 | <0.01 | *** |
| Physics - Agora Center = 0 | -0.19951 | 0.01918 | -10.402 | <0.01 | *** |
| Chemistry - Agora Center = 0 | -0.19271 | 0.019355 | -9.957 | <0.01 | *** |
| Business & Economics - Agora Center = 0 | -0.19364 | 0.0201 | -9.634 | <0.01 | *** |
| Music, Arts & Culture - Agora Center = 0 | -0.19812 | 0.02064 | -9.599 | <0.01 | *** |
| Mathematics & Statistics - Bio-environmental = 0 | 0.107356 | 0.011371 | 9.441 | <0.01 | *** |
| Information Technology - Agora Center = 0 | -0.18096 | 0.019407 | -9.325 | <0.01 | *** |
| Physics - Education = 0 | -0.07757 | 0.008358 | -9.281 | <0.01 | *** |
| Psychology - Agora Center = 0 | -0.18006 | 0.019557 | -9.207 | <0.01 | *** |
| Sport & Health - Agora Center = 0 | -0.17627 | 0.019301 | -9.132 | <0.01 | *** |
| Chemistry - Bio-environmental = 0 | -0.04215 | 0.005122 | -8.229 | <0.01 | *** |
| Education - Chemistry = 0 | 0.070767 | 0.008743 | 8.094 | <0.01 | *** |
| Bio-environmental - Agora Center = 0 | -0.15056 | 0.019415 | -7.755 | <0.01 | *** |
| Physics - Language & Communication = 0 | -0.09606 | 0.013465 | -7.134 | <0.01 | *** |
| Education - Business & Economics = 0 | 0.071696 | 0.010263 | 6.986 | <0.01 | *** |
| Music, Arts & Culture - Education = 0 | -0.07617 | 0.011284 | -6.75 | <0.01 | *** |
| Information Technology - Education = 0 | -0.05902 | 0.008858 | -6.662 | <0.01 | *** |
| Teacher Education - Agora Center = 0 | -0.14612 | 0.022292 | -6.555 | <0.01 | *** |
| Teacher Education - Mathematics & Statistics = 0 | -0.10291 | 0.015773 | -6.524 | <0.01 | *** |
| Language & Communication - Chemistry = 0 | 0.089263 | 0.013704 | 6.514 | <0.01 | *** |
| Psychology - Education = 0 | -0.05811 | 0.009153 | -6.349 | <0.01 | *** |
| Sport & Health - Education = 0 | -0.05432 | 0.008608 | -6.31 | <0.01 | *** |
| Language & Communication - Business & Economics = 0 | 0.090192 | 0.01471 | 6.132 | <0.01 | *** |
| Music, Arts & Culture - Language & Communication = 0 | -0.09467 | 0.01544 | -6.131 | <0.01 | *** |
| Social Sciences & Philosophy - Agora Center = 0 | -0.13221 | 0.022114 | -5.979 | <0.01 | *** |
| Sport & Health - Physics = 0 | 0.02325 | 0.0039 | 5.962 | <0.01 | *** |
| Education - Agora Center = 0 | -0.12195 | 0.020668 | -5.9 | <0.01 | *** |
| Mathematics & Statistics - Education = 0 | 0.078738 | 0.013397 | 5.877 | <0.01 | *** |
| Social Sciences & Philosophy - Physics = 0 | 0.067301 | 0.011473 | 5.866 | <0.01 | *** |
| Business & Economics - Bio-environmental = 0 | -0.04308 | 0.007427 | -5.8 | <0.01 | *** |
| Social Sciences & Philosophy - Mathematics & Statistics = 0 | -0.08901 | 0.015526 | -5.733 | <0.01 | *** |
| Information Technology - Bio-environmental = 0 | -0.0304 | 0.005316 | -5.719 | <0.01 | *** |
| Language & Communication - Information Technology = 0 | 0.077513 | 0.013778 | 5.626 | <0.01 | *** |
| Psychology - Language & Communication = 0 | -0.07661 | 0.013957 | -5.489 | <0.01 | *** |
| Physics - Applied Linguistics = 0 | -0.11684 | 0.021351 | -5.472 | <0.01 | *** |
| Music, Arts & Culture - Bio-environmental = 0 | -0.04755 | 0.008784 | -5.414 | <0.01 | *** |
| Sport & Health - Language & Communication = 0 | -0.07281 | 0.013612 | -5.349 | <0.01 | *** |
| Sport & Health - Bio-environmental = 0 | -0.0257 | 0.00489 | -5.256 | <0.01 | *** |
| Social Sciences & Philosophy - Chemistry = 0 | 0.060499 | 0.011753 | 5.147 | <0.01 | *** |
| Chemistry - Applied Linguistics = 0 | -0.11003 | 0.021504 | -5.117 | <0.01 | *** |
| Music, Arts & Culture - Applied Linguistics = 0 | -0.11544 | 0.022653 | -5.096 | <0.01 | *** |
| Psychology - Bio-environmental = 0 | -0.0295 | 0.005797 | -5.088 | <0.01 | *** |
| Business & Economics - Applied Linguistics = 0 | -0.11096 | 0.022162 | -5.007 | <0.01 | *** |
| Social Sciences & Philosophy - Music, Arts & Culture = 0 | 0.065903 | 0.01374 | 4.796 | <0.01 | *** |
| Social Sciences & Philosophy - Business & Economics = 0 | 0.061428 | 0.012914 | 4.757 | <0.01 | *** |
| Information Technology - Applied Linguistics = 0 | -0.09828 | 0.021551 | -4.561 | <0.01 | *** |
| Teacher Education - Physics = 0 | 0.053398 | 0.01181 | 4.522 | <0.01 | *** |
| Psychology - Applied Linguistics = 0 | -0.09738 | 0.02167 | -4.494 | <0.01 | *** |
| Language & Communication - Agora Center = 0 | -0.10345 | 0.02321 | -4.457 | <0.01 | *** |
| Sport & Health - Applied Linguistics = 0 | -0.09359 | 0.021447 | -4.364 | <0.01 | *** |
| Physics - Information Technology = 0 | -0.01855 | 0.004399 | -4.218 | <0.01 | ** |
| Social Sciences & Philosophy - Information Technology = 0 | 0.048749 | 0.011839 | 4.118 | <0.01 | ** |
| History & Ethnology - Agora Center = 0 | -0.13307 | 0.033447 | -3.979 | <0.01 | ** |
| Social Sciences & Philosophy - Psychology = 0 | 0.047846 | 0.012051 | 3.97 | <0.01 | ** |
| Psychology - Physics = 0 | 0.019455 | 0.00501 | 3.884 | <0.01 | ** |
| Teacher Education - Chemistry = 0 | 0.046596 | 0.012081 | 3.857 | <0.01 | ** |
| Sport & Health - Social Sciences & Philosophy = 0 | -0.04405 | 0.011648 | -3.782 | 0.0106 | * |
| Teacher Education - Music, Arts & Culture = 0 | 0.052 | 0.014017 | 3.71 | 0.0131 | * |
| Teacher Education - Business & Economics = 0 | 0.047525 | 0.013208 | 3.598 | 0.0197 | * |
| Mathematics & Statistics - Language & Communication = 0 | 0.060243 | 0.017049 | 3.534 | 0.0236 | * |
| Sport & Health - Chemistry = 0 | 0.016448 | 0.004672 | 3.52 | 0.0264 | * |
| Language & Communication - Bio-environmental = 0 | 0.047114 | 0.01377 | 3.421 | 0.0342 | * |
| Education - Bio-environmental = 0 | 0.028618 | 0.008858 | 3.231 | 0.0652 | . |
| Bio-environmental - Applied Linguistics = 0 | -0.06788 | 0.021549 | -3.15 | 0.0822 | . |
| Mathematics & Statistics - History & Ethnology = 0 | 0.089864 | 0.029509 | 3.045 | 0.1128 |  |
| Applied Linguistics - Agora Center = 0 | -0.08268 | 0.028516 | -2.899 | 0.1624 |  |
| Teacher Education - Information Technology = 0 | 0.034846 | 0.012165 | 2.865 | 0.1807 |  |
| Teacher Education - Psychology = 0 | 0.033943 | 0.012364 | 2.745 | 0.2374 |  |
| Teacher Education - Applied Linguistics = 0 | -0.06344 | 0.024158 | -2.626 | 0.3051 |  |
| Sport & Health - Music, Arts & Culture = 0 | 0.021852 | 0.008532 | 2.561 | 0.35 |  |
| Teacher Education - Sport & Health = 0 | 0.030148 | 0.011975 | 2.518 | 0.3814 |  |
| Sport & Health - Business & Economics = 0 | 0.017377 | 0.007127 | 2.438 | 0.4364 |  |
| Teacher Education - Language & Communication = 0 | -0.04267 | 0.01757 | -2.428 | 0.4455 |  |
| Physics - History & Ethnology = 0 | -0.06644 | 0.027592 | -2.408 | 0.4599 |  |
| Information Technology - Chemistry = 0 | 0.01175 | 0.005103 | 2.303 | 0.5412 |  |
| Music, Arts & Culture - History & Ethnology = 0 | -0.06504 | 0.028611 | -2.273 | 0.5645 |  |
| Psychology - Chemistry = 0 | 0.012653 | 0.005626 | 2.249 | 0.5844 |  |
| History & Ethnology - Chemistry = 0 | 0.059641 | 0.02771 | 2.152 | 0.6581 |  |
| History & Ethnology - Business & Economics = 0 | 0.06057 | 0.028224 | 2.146 | 0.6621 |  |
| Social Sciences & Philosophy - Applied Linguistics = 0 | -0.04953 | 0.023999 | -2.064 | 0.7228 |  |
| Psychology - Music, Arts & Culture = 0 | 0.018057 | 0.009079 | 1.989 | 0.7762 |  |
| Mathematics & Statistics - Agora Center = 0 | -0.04321 | 0.02186 | -1.977 | 0.7823 |  |
| Music, Arts & Culture - Information Technology = 0 | -0.01715 | 0.008786 | -1.952 | 0.7975 |  |
| Psychology - Business & Economics = 0 | 0.013582 | 0.007773 | 1.747 | 0.9048 |  |
| Education - Applied Linguistics = 0 | -0.03927 | 0.022682 | -1.731 | 0.9105 |  |
| Information Technology - History & Ethnology = 0 | -0.04789 | 0.027747 | -1.726 | 0.9124 |  |
| Teacher Education - Education = 0 | -0.02417 | 0.014066 | -1.718 | 0.9151 |  |
| Information Technology - Business & Economics = 0 | 0.012679 | 0.007431 | 1.706 | 0.9195 |  |
| Psychology - History & Ethnology = 0 | -0.04699 | 0.027839 | -1.688 | 0.9264 |  |
| Mathematics & Statistics - Applied Linguistics = 0 | 0.039472 | 0.023777 | 1.66 | 0.9357 |  |
| Social Sciences & Philosophy - Language & Communication = 0 | -0.02876 | 0.017352 | -1.658 | 0.9357 |  |
| Physics - Chemistry = 0 | -0.0068 | 0.004163 | -1.634 | 0.9428 |  |
| Sport & Health - History & Ethnology = 0 | -0.04319 | 0.027666 | -1.561 | 0.9609 |  |
| Social Sciences & Philosophy - Bio-environmental = 0 | 0.01835 | 0.011832 | 1.551 | 0.9631 |  |
| History & Ethnology - Applied Linguistics = 0 | -0.05039 | 0.034724 | -1.451 | 0.9797 |  |
| Language & Communication - Education = 0 | 0.018496 | 0.015484 | 1.194 | 0.9972 |  |
| Language & Communication - History & Ethnology = 0 | 0.029622 | 0.030511 | 0.971 | 0.9997 |  |
| Sport & Health - Information Technology = 0 | 0.004698 | 0.004884 | 0.962 | 0.9998 |  |
| Teacher Education - Social Sciences & Philosophy = 0 | -0.0139 | 0.016098 | -0.864 | 0.9999 |  |
| Physics - Business & Economics = 0 | -0.00587 | 0.006829 | -0.86 | 0.9999 |  |
| Language & Communication - Applied Linguistics = 0 | -0.02077 | 0.02501 | -0.831 | 1 |  |
| Social Sciences & Philosophy - Education = 0 | -0.01027 | 0.01379 | -0.745 | 1 |  |
| Sport & Health - Psychology = 0 | 0.003795 | 0.005409 | 0.702 | 1 |  |
| History & Ethnology - Bio-environmental = 0 | 0.017492 | 0.027745 | 0.63 | 1 |  |
| Music, Arts & Culture - Chemistry = 0 | -0.0054 | 0.00867 | -0.623 | 1 |  |
| Music, Arts & Culture - Business & Economics = 0 | -0.00447 | 0.010197 | -0.439 | 1 |  |
| Teacher Education - History & Ethnology = 0 | -0.01304 | 0.029816 | -0.438 | 1 |  |
| History & Ethnology - Education = 0 | -0.01113 | 0.028634 | -0.389 | 1 |  |
| Teacher Education - Bio-environmental = 0 | 0.004447 | 0.012154 | 0.366 | 1 |  |
| Physics - Music, Arts & Culture = 0 | -0.0014 | 0.008282 | -0.169 | 1 |  |
| Psychology - Information Technology = 0 | 0.000903 | 0.005804 | 0.156 | 1 |  |
| Chemistry - Business & Economics = 0 | 0.000929 | 0.007293 | 0.127 | 1 |  |
| Social Sciences & Philosophy - History & Ethnology = 0 | 0.000858 | 0.029687 | 0.029 | 1 |  |

**Supplementary Table S2:** Pairwise Tukey post-hoc test for difference between academic departments at JYU, in intercept for citation rate on CPR, in decreasing order of absolute z-value. Bonferroni-adjusted p-values reported.

| **Linear Hypotheses: intercept for the effect of citation rate on CPR is equal between two departments** | **Estimate** | **Std. Error** | **z value** | **p** |  |
| --- | --- | --- | --- | --- | --- |
| Sport & Health - Physics = 0 | -0.23978 | 0.014269 | -16.804 | <0.01 | *** |
| Physics - Agora Center = 0 | 0.969833 | 0.065551 | 14.795 | <0.01 | *** |
| History & Ethnology - Agora Center = 0 | 1.133103 | 0.078111 | 14.506 | <0.01 | *** |
| Psychology - Physics = 0 | -0.25793 | 0.018043 | -14.296 | <0.01 | *** |
| Mathematics & Statistics - Agora Center = 0 | 0.92178 | 0.067754 | 13.605 | <0.01 | *** |
| Chemistry - Agora Center = 0 | 0.896928 | 0.066073 | 13.575 | <0.01 | *** |
| Social Sciences & Philosophy - Agora Center = 0 | 0.908604 | 0.068358 | 13.292 | <0.01 | *** |
| Physics - Bio-environmental = 0 | 0.202459 | 0.015314 | 13.22 | <0.01 | *** |
| Information Technology - Agora Center = 0 | 0.868252 | 0.065776 | 13.2 | <0.01 | *** |
| Music, Arts & Culture - Agora Center = 0 | 0.873897 | 0.070663 | 12.367 | <0.01 | *** |
| Language & Communication - Agora Center = 0 | 0.846971 | 0.069766 | 12.14 | <0.01 | *** |
| Business & Economics - Agora Center = 0 | 0.797483 | 0.068187 | 11.696 | <0.01 | *** |
| Education - Agora Center = 0 | 0.790036 | 0.067739 | 11.663 | <0.01 | *** |
| Bio-environmental - Agora Center = 0 | 0.767373 | 0.066001 | 11.627 | <0.01 | *** |
| Sport & Health - Agora Center = 0 | 0.730058 | 0.065766 | 11.101 | <0.01 | *** |
| Teacher Education - Agora Center = 0 | 0.778527 | 0.071155 | 10.941 | <0.01 | *** |
| Psychology - Agora Center = 0 | 0.711903 | 0.066681 | 10.676 | <0.01 | *** |
| Applied Linguistics - Agora Center = 0 | 0.81956 | 0.080012 | 10.243 | <0.01 | *** |
| Sport & Health - Chemistry = 0 | -0.16687 | 0.016504 | -10.111 | <0.01 | *** |
| Psychology - Chemistry = 0 | -0.18503 | 0.019847 | -9.322 | <0.01 | *** |
| Psychology - History & Ethnology = 0 | -0.4212 | 0.046112 | -9.134 | <0.01 | *** |
| Sport & Health - Information Technology = 0 | -0.1382 | 0.015263 | -9.054 | <0.01 | *** |
| Sport & Health - History & Ethnology = 0 | -0.40305 | 0.044792 | -8.998 | <0.01 | *** |
| Sport & Health - Mathematics & Statistics = 0 | -0.19172 | 0.02228 | -8.605 | <0.01 | *** |
| Psychology - Mathematics & Statistics = 0 | -0.20988 | 0.024835 | -8.451 | <0.01 | *** |
| Physics - Education = 0 | 0.179797 | 0.021619 | 8.317 | <0.01 | *** |
| Psychology - Information Technology = 0 | -0.15635 | 0.01881 | -8.312 | <0.01 | *** |
| History & Ethnology - Bio-environmental = 0 | 0.36573 | 0.045131 | 8.104 | <0.01 | *** |
| Physics - Business & Economics = 0 | 0.172349 | 0.022982 | 7.499 | <0.01 | *** |
| Social Sciences & Philosophy - Psychology = 0 | 0.1967 | 0.026427 | 7.443 | <0.01 | *** |
| Chemistry - Bio-environmental = 0 | 0.129555 | 0.017414 | 7.44 | <0.01 | *** |
| Sport & Health - Social Sciences & Philosophy = 0 | -0.17855 | 0.024053 | -7.423 | <0.01 | *** |
| History & Ethnology - Education = 0 | 0.343067 | 0.047623 | 7.204 | <0.01 | *** |
| Physics - Information Technology = 0 | 0.10158 | 0.014299 | 7.104 | <0.01 | *** |
| History & Ethnology - Business & Economics = 0 | 0.33562 | 0.048256 | 6.955 | <0.01 | *** |
| Teacher Education - History & Ethnology = 0 | -0.35458 | 0.052352 | -6.773 | <0.01 | *** |
| Mathematics & Statistics - Bio-environmental = 0 | 0.154407 | 0.022957 | 6.726 | <0.01 | *** |
| Teacher Education - Physics = 0 | -0.19131 | 0.0307 | -6.231 | <0.01 | *** |
| Information Technology - Bio-environmental = 0 | 0.100879 | 0.016238 | 6.212 | <0.01 | *** |
| Information Technology - History & Ethnology = 0 | -0.26485 | 0.044795 | -5.912 | <0.01 | *** |
| Social Sciences & Philosophy - Bio-environmental = 0 | 0.14123 | 0.024679 | 5.723 | <0.01 | *** |
| Language & Communication - History & Ethnology = 0 | -0.28613 | 0.050447 | -5.672 | <0.01 | *** |
| History & Ethnology - Chemistry = 0 | 0.236175 | 0.045246 | 5.22 | <0.01 | *** |
| Psychology - Music, Arts & Culture = 0 | -0.16199 | 0.031936 | -5.073 | <0.01 | *** |
| Music, Arts & Culture - History & Ethnology = 0 | -0.25921 | 0.051695 | -5.014 | <0.01 | *** |
| History & Ethnology - Applied Linguistics = 0 | 0.313543 | 0.063879 | 4.908 | <0.01 | *** |
| Sport & Health - Music, Arts & Culture = 0 | -0.14384 | 0.029991 | -4.796 | <0.01 | *** |
| Mathematics & Statistics - Education = 0 | 0.131744 | 0.02754 | 4.784 | <0.01 | *** |
| Physics - Chemistry = 0 | 0.072905 | 0.015584 | 4.678 | <0.01 | *** |
| Social Sciences & Philosophy - History & Ethnology = 0 | -0.2245 | 0.04848 | -4.631 | <0.01 | *** |
| Education - Chemistry = 0 | -0.10689 | 0.023146 | -4.618 | <0.01 | *** |
| Psychology - Language & Communication = 0 | -0.13507 | 0.029881 | -4.52 | <0.01 | *** |
| Physics - Language & Communication = 0 | 0.122861 | 0.02732 | 4.497 | <0.01 | *** |
| Mathematics & Statistics - History & Ethnology = 0 | -0.21132 | 0.047634 | -4.436 | <0.01 | *** |
| Mathematics & Statistics - Business & Economics = 0 | 0.124297 | 0.028622 | 4.343 | <0.01 | ** |
| Sport & Health - Language & Communication = 0 | -0.11691 | 0.027806 | -4.205 | <0.01 | ** |
| Social Sciences & Philosophy - Education = 0 | 0.118568 | 0.028984 | 4.091 | <0.01 | ** |
| Teacher Education - Mathematics & Statistics = 0 | -0.14325 | 0.035096 | -4.082 | <0.01 | ** |
| Chemistry - Business & Economics = 0 | 0.099445 | 0.024424 | 4.072 | <0.01 | ** |
| Teacher Education - Chemistry = 0 | -0.1184 | 0.031787 | -3.725 | 0.014 | * |
| Social Sciences & Philosophy - Business & Economics = 0 | 0.11112 | 0.030013 | 3.702 | 0.0156 | * |
| Physics - History & Ethnology = 0 | -0.16327 | 0.044486 | -3.67 | 0.0165 | * |
| Teacher Education - Social Sciences & Philosophy = 0 | -0.13008 | 0.036231 | -3.59 | 0.023 | * |
| Information Technology - Education = 0 | 0.078217 | 0.022262 | 3.513 | 0.0287 | * |
| Music, Arts & Culture - Bio-environmental = 0 | 0.106524 | 0.030498 | 3.493 | 0.0308 | * |
| Psychology - Business & Economics = 0 | -0.08558 | 0.026003 | -3.291 | 0.06 | . |
| Physics - Music, Arts & Culture = 0 | 0.095935 | 0.029524 | 3.249 | 0.0682 | . |
| Psychology - Education = 0 | -0.07813 | 0.024807 | -3.15 | 0.0892 | . |
| Physics - Applied Linguistics = 0 | 0.150272 | 0.047761 | 3.146 | 0.0933 | . |
| Information Technology - Business & Economics = 0 | 0.070769 | 0.023587 | 3 | 0.1362 |  |
| Teacher Education - Information Technology = 0 | -0.08973 | 0.031136 | -2.882 | 0.186 |  |
| Sport & Health - Business & Economics = 0 | -0.06743 | 0.023575 | -2.86 | 0.195 |  |
| Psychology - Bio-environmental = 0 | -0.05547 | 0.019592 | -2.831 | 0.2079 |  |
| Language & Communication - Bio-environmental = 0 | 0.079598 | 0.028349 | 2.808 | 0.22 |  |
| Sport & Health - Education = 0 | -0.05998 | 0.022249 | -2.696 | 0.2841 |  |
| Social Sciences & Philosophy - Physics = 0 | -0.06123 | 0.023486 | -2.607 | 0.3393 |  |
| Music, Arts & Culture - Education = 0 | 0.083862 | 0.034083 | 2.461 | 0.4433 |  |
| Mathematics & Statistics - Information Technology = 0 | 0.053528 | 0.022291 | 2.401 | 0.4869 |  |
| Teacher Education - Music, Arts & Culture = 0 | -0.09537 | 0.040435 | -2.359 | 0.5209 |  |
| Mathematics & Statistics - Language & Communication = 0 | 0.074809 | 0.032185 | 2.324 | 0.5477 |  |
| Sport & Health - Bio-environmental = 0 | -0.03732 | 0.016216 | -2.301 | 0.5643 |  |
| Physics - Mathematics & Statistics = 0 | 0.048052 | 0.021647 | 2.22 | 0.6286 |  |
| Music, Arts & Culture - Business & Economics = 0 | 0.076414 | 0.034963 | 2.186 | 0.6561 |  |
| Psychology - Applied Linguistics = 0 | -0.10766 | 0.049277 | -2.185 | 0.6562 |  |
| Teacher Education - Psychology = 0 | 0.066624 | 0.032996 | 2.019 | 0.7734 |  |
| Mathematics & Statistics - Applied Linguistics = 0 | 0.10222 | 0.050708 | 2.016 | 0.7746 |  |
| Sport & Health - Applied Linguistics = 0 | -0.0895 | 0.048044 | -1.863 | 0.862 |  |
| Social Sciences & Philosophy - Language & Communication = 0 | 0.061632 | 0.03342 | 1.844 | 0.8714 |  |
| Language & Communication - Education = 0 | 0.056935 | 0.032165 | 1.77 | 0.9042 |  |
| Teacher Education - Language & Communication = 0 | -0.06844 | 0.038822 | -1.763 | 0.9077 |  |
| Language & Communication - Chemistry = 0 | -0.04996 | 0.028537 | -1.751 | 0.9121 |  |
| Information Technology - Chemistry = 0 | -0.02868 | 0.016529 | -1.735 | 0.9185 |  |
| Social Sciences & Philosophy - Applied Linguistics = 0 | 0.089043 | 0.051503 | 1.729 | 0.9206 |  |
| Social Sciences & Philosophy - Information Technology = 0 | 0.040351 | 0.024061 | 1.677 | 0.9366 |  |
| Chemistry - Applied Linguistics = 0 | 0.077368 | 0.048469 | 1.596 | 0.9584 |  |
| Teacher Education - Sport & Health = 0 | 0.04847 | 0.03113 | 1.557 | 0.9665 |  |
| Language & Communication - Business & Economics = 0 | 0.049488 | 0.033096 | 1.495 | 0.9766 |  |
| Music, Arts & Culture - Mathematics & Statistics = 0 | -0.04788 | 0.034101 | -1.404 | 0.987 |  |
| Business & Economics - Bio-environmental = 0 | 0.03011 | 0.024216 | 1.243 | 0.9963 |  |
| Bio-environmental - Applied Linguistics = 0 | -0.05219 | 0.048361 | -1.079 | 0.9992 |  |
| Mathematics & Statistics - Chemistry = 0 | 0.024852 | 0.023174 | 1.072 | 0.9993 |  |
| Information Technology - Applied Linguistics = 0 | 0.048692 | 0.048049 | 1.013 | 0.9996 |  |
| Music, Arts & Culture - Applied Linguistics = 0 | 0.054337 | 0.054539 | 0.996 | 0.9997 |  |
| Education - Bio-environmental = 0 | 0.022663 | 0.022927 | 0.988 | 0.9997 |  |
| Social Sciences & Philosophy - Music, Arts & Culture = 0 | 0.034706 | 0.035278 | 0.984 | 0.9998 |  |
| Sport & Health - Psychology = 0 | 0.018154 | 0.018793 | 0.966 | 0.9998 |  |
| Language & Communication - Information Technology = 0 | -0.02128 | 0.027813 | -0.765 | 1 |  |
| Music, Arts & Culture - Chemistry = 0 | -0.02303 | 0.030661 | -0.751 | 1 |  |
| Teacher Education - Applied Linguistics = 0 | -0.04103 | 0.055162 | -0.744 | 1 |  |
| Music, Arts & Culture - Language & Communication = 0 | 0.026926 | 0.037935 | 0.71 | 1 |  |
| Education - Applied Linguistics = 0 | -0.02952 | 0.050695 | -0.582 | 1 |  |
| Teacher Education - Business & Economics = 0 | -0.01896 | 0.035933 | -0.528 | 1 |  |
| Language & Communication - Applied Linguistics = 0 | 0.027411 | 0.053358 | 0.514 | 1 |  |
| Social Sciences & Philosophy - Chemistry = 0 | 0.011676 | 0.024893 | 0.469 | 1 |  |
| Social Sciences & Philosophy - Mathematics & Statistics = 0 | -0.01318 | 0.029005 | -0.454 | 1 |  |
| Business & Economics - Applied Linguistics = 0 | -0.02208 | 0.05129 | -0.43 | 1 |  |
| Teacher Education - Bio-environmental = 0 | 0.011154 | 0.031616 | 0.353 | 1 |  |
| Teacher Education - Education = 0 | -0.01151 | 0.035077 | -0.328 | 1 |  |
| Education - Business & Economics = 0 | -0.00745 | 0.028599 | -0.26 | 1 |  |
| Music, Arts & Culture - Information Technology = 0 | 0.005645 | 0.03 | 0.188 | 1 |  |

## Footnotes

1 Note on terminology: we use “scholarly” and “output” here as umbrella terms to include all serious academic study and different formats and channels of its communication, some of which might be seen to be excluded by terms “scientific” and “article” which we use here only when referring to earlier work by others using those terms.

## Notes

https://gitlab.jyu.fi/jyucite/published_cpr

